## Supplementary file 1 for "CPT1A Mediates Radiation Sensitivity in Colorectal Cancer"

**Supplementary file 1. The ‾D0 and N value of multi-target, single-hit model in all cell lines.**

| Cell line | HCT-15 | RKO | HCT 116 | HT-29 | Caco-2 | SW480 | SW620 |
| --- | --- | --- | --- | --- | --- | --- | --- |
| ‾D_0_ | 3.929 | 1.808 | 1.902 | 4.820 | 2.423 | 2.752 | 2.558 |
| N | 1.267 | 1.698 | 1.162 | 2.327 | 1.816 | 1.518 | 1.058 |
