## Supplementary file 2 for "CPT1A Mediates Radiation Sensitivity in Colorectal Cancer"

**Supplementary file 2. Primers used for the real-time PCR**

| Gene symbol | Forward Primer | Reverse Primer |
| --- | --- | --- |
| β-actin | CATGTACGTTGCTATCCAGGC | CTCCTTAATGTCACGCACGAT |
| CPT1A | ATCAATCGGACTCTGGAAACGG | TCAGGGAGTAGCGCATGGT |
| PPARA | TTCGCAATCCATCGGCGAG | CCACAGGATAAGTCACCGAGG |
| PPARG | GGGATCAGCTCCGTGGATCT | TGCACTTTGGTACTCTTGAAGTT |
| PGC1A | GCTTTCTGGGTGGACTCAAGT | GAGGGCAATCCGTCTTCATCC |
| FOXM1 | ATACGTGGATTGAGGACCACT | TCCAATGTCAAGTAGCGGTTG |
| SOD1 | GGTGGGCCAAAGGATGAAGAG | CCACAAGCCAAACGACTTCC |
| SOD2 | GGAAGCCATCAAACGTGACTT | CCCGTTCCTTATTGAAACCAAGC |
| SOD3 | ATGCTGGCGCTACTGTGTTC | CTCCGCCGAGTCAGAGTTG |
| CAT | TGGAGCTGGTAACCCAGTAGG | CCTTTGCCTTGGAGTATTTGGTA |
